# Tree canopy influences ground level atmospheric electrical and biogeochemical variability

**DOI:** 10.1101/2021.03.17.435822

**Authors:** Ellard R. Hunting, Sam J. England, Daniel Robert

**Author notes:** Correspondence: Ellard R. Hunting, Daniel Robert.

## Abstract

Static electric fields in the atmosphere are increasingly recognized to interact with various organisms over several levels of biological organization. Recently, a link between atmospheric electrical variations and biogeochemical processes has been established in the context of open fields, yet biological structures like trees produce substantial alterations in atmospheric electric properties. Here, we assess whether these structural changes affect the dynamics of both biogenic and abiotic electrical landscapes and their relation to geochemical processes. To this end, we theoretically assess how trees alter their surrounding electric fields and empirically compare the temporal dynamics of atmospheric potential gradients, positive ions in the near-ground level atmosphere and soil electrochemical properties in an open field and under a tree. The developed model of electric fields around trees provides insight into the extent to which trees shield underlying electric landscape, revealing that a substantial increase in atmospheric potential gradient only marginally affects the electric field under the canopy. We further show that soil electrochemical properties are tied to temporal dynamics of positive ions in the near-ground level atmosphere, and that the presence of a tree reduces the temporal variability in both ground level positive ions concentrations and soil redox potential. This suggests that a tree can have a stabilizing effect on drivers of temporal variability in atmospheric electricity and soil electro-chemistry, thereby likely indirectly influencing soil microorganisms and processes as well as electro-sensitive organisms that perceive and utilize atmospheric electric fields.

## Introduction

The atmosphere is host to various electrical phenomena, spanning immense dimensions that range from the local production of single electrons and ions to the global electric circuit (Rycroft et al., 2008). Evidence is now emerging that static electric fields in the atmosphere can interact with various organisms over several levels of biological organization (e.g., molecules, cells, organisms, communities) (Clarke et al., 2013; Greggers et al., 2013; Morley et al., 2018; Hunting et al., 2020). These electric fields – as well as their dynamics – have also been linked to biogeochemical processes below the Earth’s surface by means of a charge separation between the negatively charged Earth’s interior and positive charge sources in the atmosphere (Hunting et al., 2019). These atmospheric sources range from a ground level build-up of positive ions (Adkins, 1959; Crozier, 1965) to positively charged fair weather regions that are fuelled by thunderstorm regions around the globe as part of the global electric circuit (Wilson, 1903; Rycroft et al., 2008; Fdez-Arroyabe et al., 2021). Consequently, variations in atmospheric electrical fields and charge flow can be observed to operate at both local and global spatial scales. Electric fields caused by these charge separations have been observed to drive a subsurface migration of nutrients essential to microbial metabolism, thereby affecting microbial processes in both soils and aquatic sediments (Hunting et al., 2019).

The conceptual and empirical foundation underlying the link between atmospheric electrical variations and biogeochemical processes has been established in the context of open fields (e.g., grasslands) and freshwater systems. However, both empirical data and mathematical models have shown that objects like trees produce substantial alterations in the electric landscape, in particular the electric field strength in the surrounding air (Arnold et al., 1965; Williams et al., 2005; Bowker and Crenshaw 2007; Clarke et al., 2016; Morley and Robert, 2018). In effect, as positive charges in the atmosphere are drawing a negative mirror charge to the surface of the trees, such as the sharp extremities of needles, and/or other anatomical features with high curvature, trees produce larger fields than flat surfaces (low curvature) (Feynman et al. 1964; Clarke et al., 2017). Thus, as plants display some mobility in their surface electric charge carriers (electrons, charged molecules and ions), the electric field surrounding them will take up geometries that are influenced by the strength of atmospheric potential gradients as well as the plants’ height and morphology. Likewise, large plants, such as trees, have been reported to contribute substantially to local variations in ground level atmospheric ions (e.g., Jayaratne et al., 2011) that are generated largely by transpiration (See Fig. 1 for a conceptual and mathematical representation of electric fields and ions around trees). It thus becomes apparent that trees and their canopies affect the electric landscape around and below them. Questions are thus arising whether these structural changes affect the dynamics of both biogenic and abiotic electrical landscapes and their relation to geochemical processes. In light of the largely uncharacterised and empirically elusive near-ground (0 to 2m) atmospheric electric properties and their unsuspected roles in geochemical changes in both near-ground and subsurface environments, we aim to contribute to the exploration of this electric ecology. To this end, we theoretically assess how trees alter their surrounding electric fields and empirically compare the temporal dynamics of atmospheric potential gradients, positive ions in the near-ground level atmosphere and soil electrochemical properties in an open field and under a tree.

**Figure 1:**
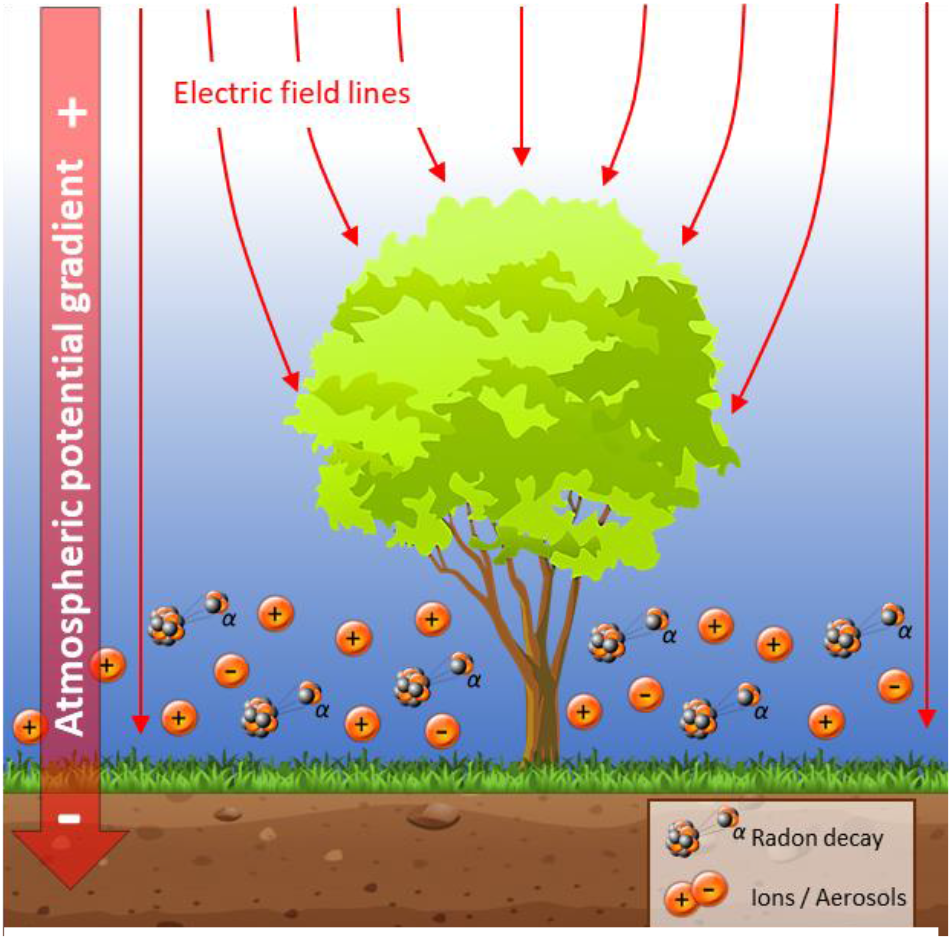
Conceptual impression of the electric landscape around trees.

## Methods

### Model of atmospheric electric fields

Since modelling approaches with two-dimensional geometries published to date are potentially prone to exaggerating the electrical shielding effect of most structures, we modelled a three-dimensional geometry to assess the effect of a tree on local atmospheric electric fields. Modelling was performed using finite element analysis within COMSOL Multiphysics® v. 5.4 (COMSOL AB, Stockholm, Sweden) utilising the “Electric Currents” interface within the “AC/DC” module. The tree-dimensional geometry consisted of a 300m × 300m × 300m cube within which the model operated. The model ground level was occupied by a 300m × 300m × 2m block of soil. At its centre, an approximate representation of a 30m tall deciduous tree, with a 2m wide trunk, and a canopy spanning roughly 30m was positioned. This tree could be regarded as a model of an oak or an alder tree. The remainder of the geometry was assigned as air. The upper surface of this air column was given an electrical potential typical of a 300m altitude in various meteorological conditions, representative of the atmospheric potential gradient (Wilson 1903; Bennett & Harrison 2007). The electrical ground was defined 2m below the upper surface of the soil (a typical groundwater table depth). Meshing of this geometry was physics-controlled, set to “extremely fine” (Supplementary Figure S1). Electrical properties of all materials were obtained from measurements or estimates from primary literature. The electrical conductivity, σ, and relative permittivity, ε_r_, were defined as: for soil, σ = 0.05 S/m, ε_r_ = 15 (Rhebergen et al., 2002; Brovelli and Cassiani, 2011); for living trees, σ = 2 × 10^−4^ S/m, ε_r_ = 12 (Gora and Yanoviak, 2014; Suojanen et al., 2001); and for air, σ = 1 × 10^−14^ S/m, ε_r_ = 1 (Hogg 1939; Higazi and Chalmers, 1966). Model outputs presented for this study were produced by plotting data from two-dimensional slices or one-dimensional cut lines through areas of interest within the three-dimensional geometry. An identical model, except with no tree present, was constructed as a reference.

### Study site and analytical techniques

Experiments were performed at the University of Bristol, School of Veterinary Sciences, Langford UK, between October 2018 and February 2019, in which measurements took place during fair weather with occassional cloud cover (condition defined in Harrison and Nicoll, 2018). Meteorological parameters where continuously measured on site using a Maximet (GMX501) weather station, and included temperature (°C), humidity (RH), air pressure (mB), solar radiation (W/m^2^), windspeed (m/s) and precipitation (mm/min). The atmospheric potential gradients were continuously measured on site using a fieldmill (Boltek EFM 100 Field Mill upside up configuration), positioned with its rotor facing up on top of a single vertical pole at 1.8m above ground. Ground was covered with short (<10cm) grass during the measurement period. Atmospheric ion concentrations were measured every second using two cylindrical capacitor-based ion-counters operating on the ‘Gerdien-tube’ principle (Alpha-lab Inc., Salt Lake City, UT, USA), calibrated in open field conditions and recorded onto a PC using a National Instruments (Austin, Texas) DAQ system. Redox potentials (Eh) in soils were measured simultaneously every 20 minutes using two permanently installed gold-plated Eh-electrodes connected to a Hypnos 3 data logger and corrected for the reference electrode and measured pH. Details on the Eh monitoring devices and electrodes are described elsewhere (Vorenhout et al., 2011).

### Experimental approach

A time-series was recorded between 13-11-2018 and 17-11-2018 in open field conditions to assess whether soil redox potential is coupled to atmospheric processes, focussing on the relation between soil redox potential, meteorological variables and the atmospheric potential gradient. A second time-series was measured between 24-10-2018 and 31-10-2018 in both open field conditions and next to the trunk of a grey alder (*Alnus incana*), focussing on the relation between soil redox potential and concentration of positive ions in the atmosphere near the ground. We used standardized soil microcosms to assure soils were comparable between both tree and open field conditions (i.e., sufficient water content, exclusion of ground water flow). To this end, 400 mL glass beakers were filled with a pre-wetted standardized soil mixture (quartz and turf, 3:1). Positive ground level atmospheric ions and soil redox potential were measured simultaneously in soil microcosms at 1 cm depth positioned at the base of a stand-alone tree and an open field, 5 meters away from the margin of the canopy of the tree.

A detailed comparative assessment between ground level atmospheric electric variations between trees and open field was based on ground level positive ions. A series of measurements in time were performed using soil microcosms as described above, but focussing on the period between 10:00 am and 20:00 pm to avoid large influences of diel fluctuations typically observed in ground level atmospheric ion concentrations (e.g. Jayaratne et al., 2011). This was repeated 6 times with freshly prepared microcosms. Since different sampling intervals do not allow for cross-correlation of different time-series, co-variability of ground level atmospheric ion concentrations and soil redox potential was visualized by plotting moving averages. Temporal variability of the obtained measurements were subsequently assessed with a Wavelet transform spectral analysis in PAST (Hammer, 2001). This form of spectral analysis proved a valuable approach in evaluating spatial and temporal patterns in ecological research by creating a composite measure of temporal variance for each treatment or spatially separated samples over time (Bradshaw and Spies, 1992; Ibarra-Junquera et al., 2006; Escalante-Minakata et al., 2009). As applied here, we followed the procedures described in detail by Hunting et al. (2015). Amplitude of temporal variation were thus characterised, allowing the comparison between simultaneous or spatially separated measurements, in which information on the specific timeframes of variability is maintained. These temporal variances were used to obtain the mean overall variance in each treatment. Finally, differences between trees and open-field variances were subsequently assessed using a small sample corrected Epps-Singleton test for equal distributions (Epps and Singleton, 1986).

## Results

Modelling using finite element analysis demonstrates that the presence of a tree has a marked impact on the surrounding electric field (Fig. 2ABC). Beneath the tree, electric field strengths are reduced by more than an order of magnitude, supporting the hypothesis that trees act as electrical shields, greatly diminishing variability in electrical conditions beneath their canopies. A horizontal transect taken through the model (Fig. 2B) shows that for a large tree in an open field, this shielding effect persists beyond the immediate vicinity of the tree, with the electric field strength continuing to be reduced in excess of 100m away from the trunk in comparison to no tree being present. In contrast to this arboreal suppression of electric fields beneath the canopy, we conversely find that the electric field strength immediately above the tree is amplified above levels expected if no tree were present. Examining the vertical transect through the model, positioned 5 meters away from the trunk (Fig. 2C), emphasises the effect of shielding, whereby significant amplification takes place near the treetop, persisting over significant distances, with some visible elevation of electric field strength even at 100m above the tree canopy. Under a fair-weather atmospheric potential gradient of 100V/m, or unstable conditions presenting 1kV/m, the electric field in the immediate vicinity of the treetop can reach 450V/m and 4.5kV/m, respectively.

**Figure 2:**
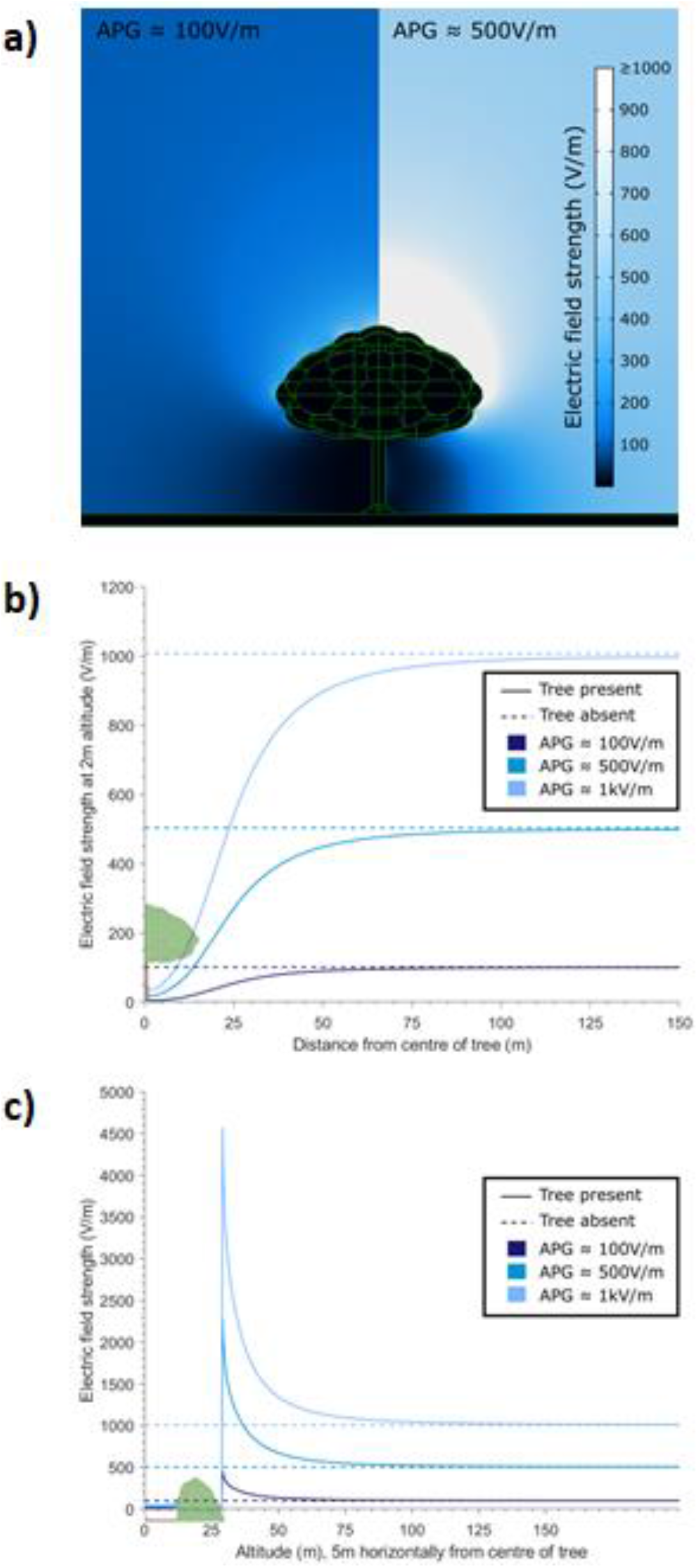
Outputs from a three-dimensional finite element analysis model of the electric fields surrounding a tree stood in open ground, exposed to various vertical atmospheric potential gradients (APGs). **(a)** two-dimensional slice through the centre of the tree in two different APG strengths, 100V/m (left) and 500V/m (right), showing the resultant electric landscape. **(b)** cutline horizontal transects taken through the model travelling from the centre of the tree outwards for 150m, at an altitude of 2m, for various APG strengths, exemplifying the electrical shielding effect of the tree both immediately underneath the tree and for large distances away from the canopy. Model outputs with a tree present (solid) are compared with identical models with no tree present (dashed) **(c)** cutline vertical transects taken through the model, travelling from the soil surface upwards for 200m, 5m horizontally from the centre of the tree, for various APG strengths, showing both the shielding effect beneath the canopy and the amplification of the APG above the tree. Again, model outputs with a tree present (solid) are compared with identical models with no tree present (dashed).

Soil redox potential did not clearly covary with any of the measured atmospheric meteorological variables (Supplementary fig. S1). An increase in redox potential and subsequent emergence of a diel cycle coincided with a period of prolonged precipitation on the 27^th^ of October, suggesting the soil was not necessarily saturated with water throughout the measurement period (e.g., Cusell et al., 2015). Diel cycles in soil redox potential between 29^th^ and 31^st^ of October 2018 did not seem to be directly governed by changes in the atmospheric potential gradient.

The 5-day measurement period between 13^th^ and 17^th^ of November 2018 shows that ground-level positive atmospheric ions and soil redox potential in soil microcosms in the open field have distinct diel cycles (Fig. 3AB). In contrast, near the trunk of a tree, soil redox potential revealed substantially (~80%) lower amplitude in diel variation, and ground-level positive atmospheric ions did not appear to have any diel rhythm (Fig. 3AB).

**Figure 3:**
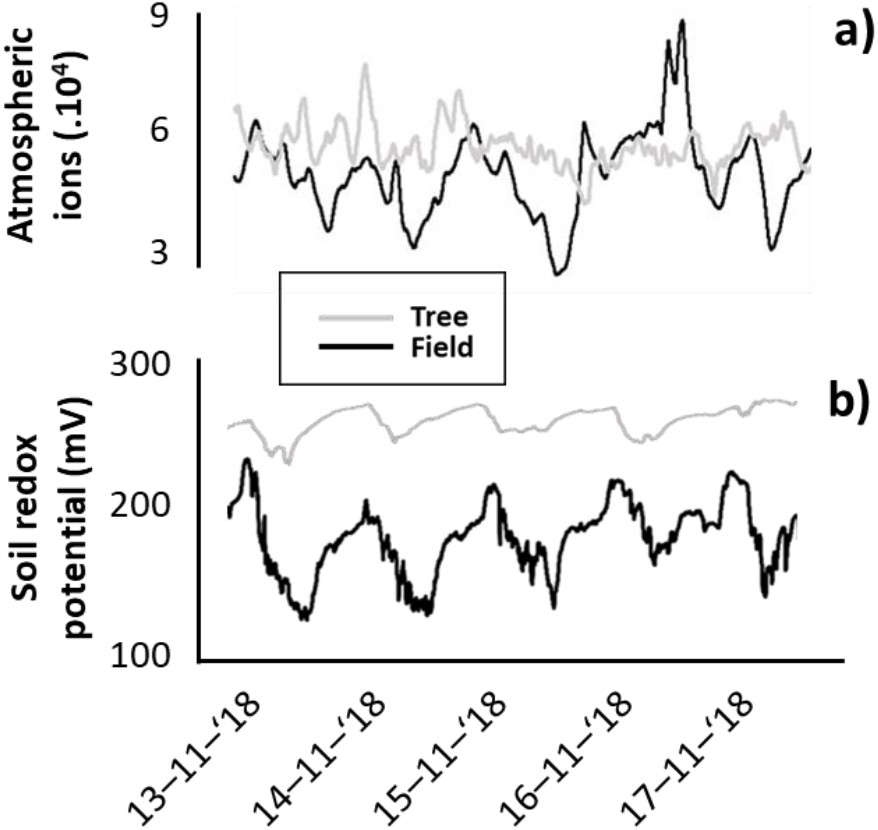
Co-variability of positive atmospheric ions and soils redox potential.

A representative example of the measured time-series similarly reveals that the moving averages of positive atmospheric ions and soils redox potential in soil microcosms covary during the day (Fig 4AB). Comparison of Wavelet variances throughout the entire measurement period across the replicated time-series (Fig. 4CD) suggests that temporal variability was lower for both ground level atmospheric ion concentrations and soil redox potential near a tree compared to Wavelet variances in the open field (Epps-Singleton-derived error: W^2^ = 63.94; *p* < 0.001 and W^2^= 1439.1; *p* < 0.001, respectively).

**Figure 4:**
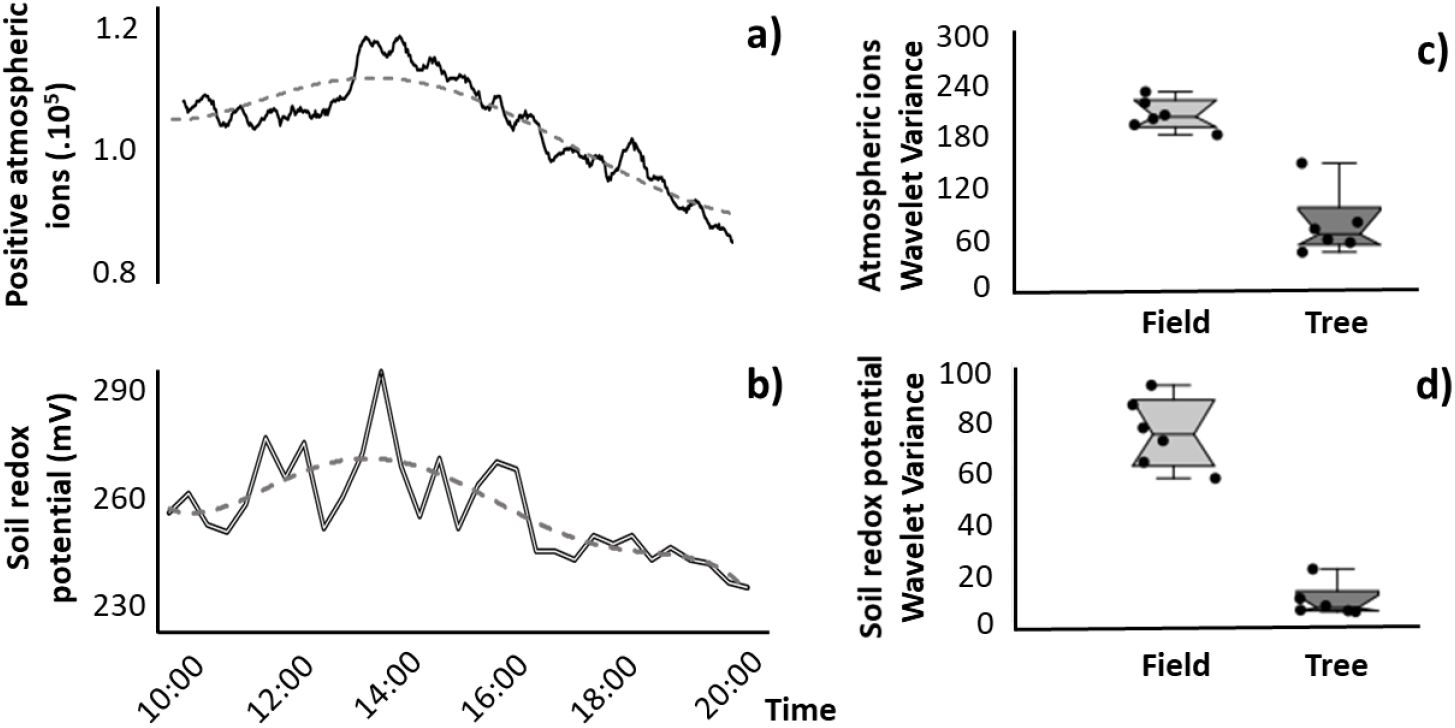
Representative example of the measured time-series revealing co-variability of A) positive atmospheric ions and B) soils redox potential in soil microcosms, in which dashed line represents the hourly moving average. Comparison of temporal variability across the replicated time-series expressed as Wavelet variances between near a tree and in an open field for C) near ground-level atmospheric positive ions and D) soil redox potential.

## Discussion

The developed model of the electric fields around trees exemplifies the drastic influence of trees and other large plants on their electrical environment, even when only considering passive electrical interactions. Undoubtedly, an even more marked effect can be expected in tree lines or forests due to a cumulative effect of collections of plants and trees (see also Williams et al., 2005). Importantly, the extension of previous two-dimensional models on electrical shielding by trees (Clarke et al., 2017; Morley and Robert, 2018) into three dimensions corroborates the influence of vegetation on their local electrical environment. The present three-dimensional model confirms that prior findings on the shielding ability of trees using a two-dimensional approach were qualitatively valid yet may not have been entirely numerically accurate. Previous studies demonstrated that tree canopies can alter atmospheric electric fields, almost nullifying them near the base of the tree trunk, while the reach of this shielding effect extended beyond the canopy cover by 5 – 20m with increasing distance from the tree (Märcz and Harrison, 2003; Williams et al., 2005; Clarke et al., 2017; Morley and Robert, 2018). Our three-dimensional model confirms that the electric field strength is reduced to near zero, and reveals that a substantial (5 to 10 fold) increase in atmospheric potential gradient only marginally affects electric fields under the tree canopy, articulating the extent to which a tree canopy shields its underlying electric landscape and distorts the atmospheric potential gradient beyond its own span. Remarkably, above the canopy the local electric field can be greatly enhanced, generating features in the electric landscape otherwise absent in open fields.

Local sources of atmospheric electricity in continental environments often remain ambiguous, relying on both globally driven atmospheric potential gradients and local sources of ionization (e.g., Wright et al., 2000). The absence of correlation between atmospheric potential gradients and other metereological parameters in our study strongly supports the notion that ground level atmospheric ion concentrations above soils are driven by local concentrations of atmospheric ions, charged aerosols and radionuclides (Kubicki etal., 2016). These ground level ion concentrations can be substantially higher around trees, observable especially in forests, compared to open fields due to transpiration of ions and radon by vegetation (e.g. Ling, et al., 2010; Jayaratne et al., 2011). Importantly, since transpiration is drastically reduced in winter, atmospheric ions between tree canopy cover and open fields are not expected to markedly differ in magnitude, suggesting that, in the present study, local sources of atmospheric ions such as soil radon exhalation and air pollution (e.g. fuel exhausts) made up the electric charges at near-ground level atmosphere in both open fields and near vegetation.

While several studies have focussed on how trees affect the local electric landscape, it remains largey unknown how trees, as large partially conductive and dielectric structures, affect the temporal dynamics of atmospheric potential gradients, and whether and how this affects multiscale chemical and biological processes. We observed clear differences in the dynamics between both ground level atmospheric ions and soil redox potential between the open field and below a tree canopy over a 5-day measuring period. Although clear diel cycles were visible in both ions and redox potential in open fields, diel cycles were not observed in atmospheric ions near the trunk of the tree. Soil redox potential was observed to have diel cycles, yet their magnitude was substantially lower (~10%) compared to open field conditions. A closer assessment of the temporal variability near a tree and the open field revealed that soil electrochemical properties are tied to temporal dynamics of positive ions in the near-ground level atmosphere, and that the presence of a tree reduced the temporal variability in both ground level positive ions concentrations and soil redox potential. This shows that, while ground level atmospheric ions were of primarily local origin, a tree can have a stabilizing effect on drivers of temporal variability. It is important to consider that various sources are known to contribute to ground-level electric variability, including windspeed, soil radon exhalation, tree transpiration and air-earth currents driven by atmospheric potential gradients (Jayaratne et al., 2011; Kubicki et al., 2016). Since our measurements were obtained in winter and near a single tree, it is unlikely that wind speed and tree transpiration were the source of variability. Likewise, soil radon exhalation is expected to add variability near trees provided tree soils are substantially more permeable compared to grasslands (Alaoui et al., 2011; Holthusen et al., 2018). Therefore, the most likely source of variability in this study is of atmospheric origin, in which a tree and its canopy appear to shield ground-level electric dynamics from atmospheric influences. This offers a plausible explanation to the frequently observed differences in the temporal characteristics of soil and sediment electrochemical signatures between open fields and near vegetation.

The observed interplay between vegetation, atmospheric electrical variations and soil redox potential is ecologically relevant since redox potential is an important driver of bacterial community structure and metabolism in both soils and aquatic sediments (Newman and Banfield, 2002; Bertics and Ziebis, 2009; Hunting and van der Geest, 2011; Hunting et al., 2012), and oscillations in redox conditions are known to change microbial community composition (Pett-Ridge and Firestone, 2005) and promote bacterial metabolic processes (Aller, 1994). These changes in microbial metabolism can be attributed to the mobilization of microbial nutrients due to changes in the physico-chemical environment (Aller, 1994) as well as the migration of respiratory ions along electrochemical gradients and an electric field (Hunting et al., 2019). Bacteria themselves have been observed to occupy an apparent “redox niche” in which they actively migrate towards the most favourable electrochemical conditions (Bespalov, 1996) or actively control the redox conditions in their immediate surroundings by membrane bound and secreted redox mediators (Hunting and Kampfraath, 2013). The observed influence of local dynamics in atmospheric electricity governed by vegetation may thus affect microbial processes in soils and aquatic sediments in vegetated areas, and likely carries wider implications for electro-sensitive organisms (e.g., pollinators, ballooning spiders, and perhaps other arthropod species) that perceive and utilize atmospheric electric fields.

## Acknowledgements

EH received financial support from the Swiss National Science Foundation, SNF (CRSK-2 190855). DR is funded by the BBSRC (grant BB/T003235/1) and the European Research Commission (ERC-ADG 743093), supporting EH and SE.

---

**Supplementary Figure S1:**
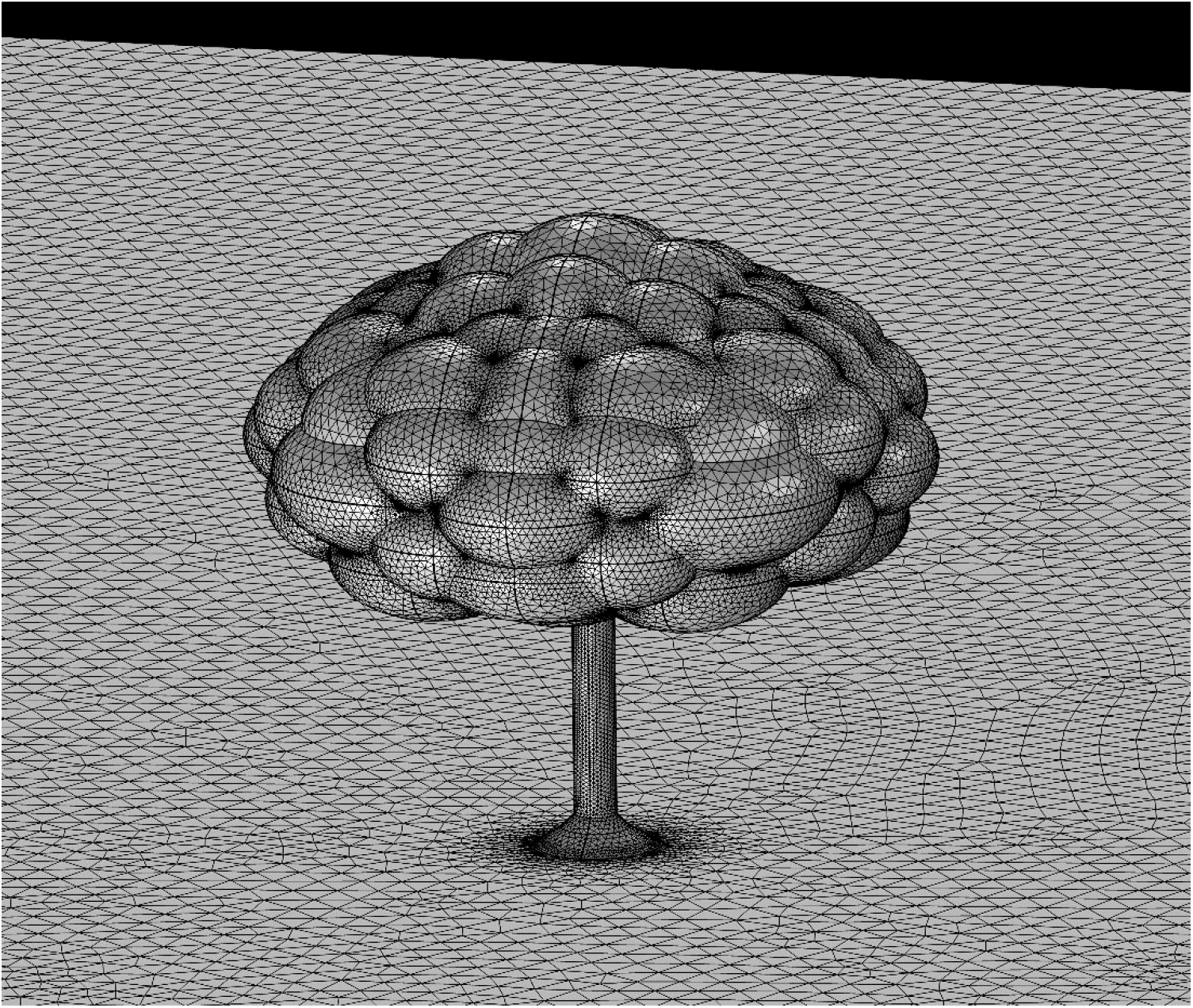
Three-dimensional geometry of the tree used for finite element analysis.

**Supplementary Figure S2:**
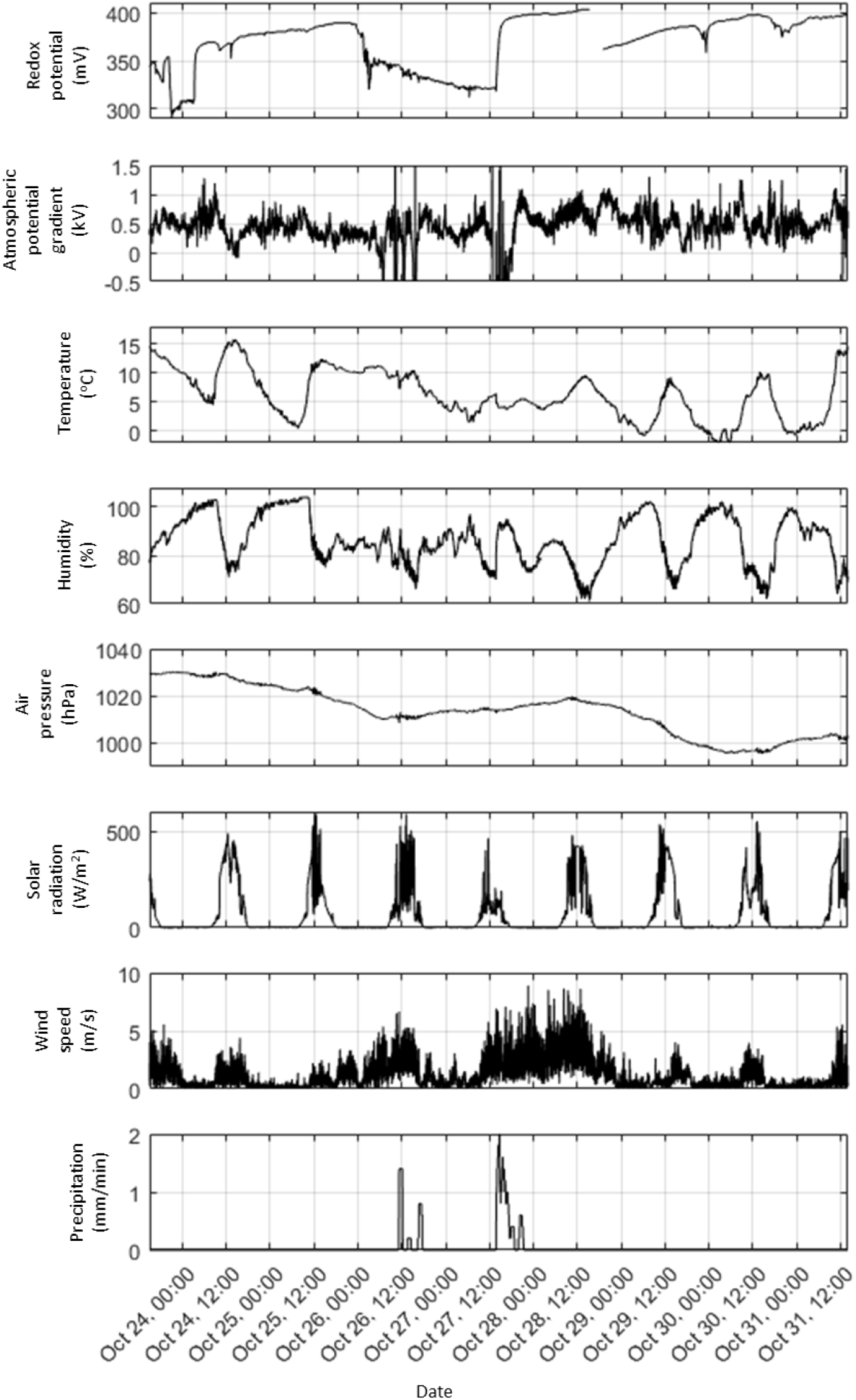
Soil redox potential and meteorological parameters at the field site.

